# Sleep Medicine Knowledge Among Graduating Medical Students in Lebanon During an Economic and Political Crisis

**DOI:** 10.1101/2021.07.09.451820

**Authors:** Raissa Aoun, Victor Zibara, Christy Costanian, Hrayr Attarian, Sola Aoun Bahous

**Affiliations:** Department of Neurology, New York University Langone Hospital, Brooklyn, NY, USA; Gilbert and Rose-Marie Chagoury, School of Medicine, Lebanese American University, Byblos, Lebanon; Department of Neurology, Northwestern Memorial Hospital, Chicago, IL, USA

**Keywords:** sleep medicine, medical education, ASKME survey, medical schools

## Abstract

**Objectives:** Sleep disorders are prevalent and underrecognized during both economic and political crises. They are a major reason for poor overall health and decreased quality of life. Sleep medicine education is limited at most medical schools, resulting in limited awareness of this important aspect of healthcare. The aim of the study is to assess sleep medicine knowledge of graduating medical students in Lebanon and to assess their readiness to tackle sleep health issues in a country during an unprecedented crisis.

**Methods:** Final-year medical students at 7 medical schools in Lebanon were invited to fill a survey between January 2020 and March 2021. The Assessment of Sleep Knowledge in Medical Education survey was used to assess their knowledge in sleep medicine. The curriculum organizers at the medical schools were also surveyed. Student’s t-test was used for analysis.

**Results:** 158 and 58 students completed the survey during 2020 and 2021, with a mean overall score on sleep knowledge of was 17.5 and 15.9 /30, respectively. There was no difference in mean knowledge scores by gender, age, American versus European medical school systems, and between medical schools that included sleep medicine in their curriculum versus those that did not.

**Conclusions:** Presence of sleep medicine education in the curriculum was associated with higher scores on ASKME among graduating Lebanese medical students. Overall, the new crop of physicians in Lebanon possesses a relatively good knowledge base in sleep medicine. Nevertheless, more effort should be made to uniformly maintain this level of sleep education.

## Introduction

Since Fall of 2019, Lebanon has been in political and economic turmoil.(1) This situation has been compounded with the start of the COVID19 global pandemic.(2) Moreover, political instability ensued because of the absence of a functioning government, rapid devaluation of the Lebanese pound, dramatic increase in poverty and random bursts of violence.(3) All these elements have been shown to have a significant impact on sleep and subsequently overall health and quality of life.(4,5) To add, poor sleep is itself associated with high risk of later trauma related distress.(6)

Even shortly before the current turmoil prevalence of chronic insomnia was high, 4.5% in a sample of the adult population of the Greater Beirut area.(7) Sleep disorders are also common among university students as well as psychiatric patients across the various diagnoses in Lebanon.(8–10) Anxiety and depression are widespread which could translate to a high prevalence of comorbid insomnia.(11) Despite this increasing burden of sleep related issues, there are no studies, to our knowledge, tackling sleep medicine knowledge among Lebanese medical students; moreover, only a few studies worldwide have assessed sleep medicine knowledge in medical students in relation to teaching and curriculum design.

Medical schools in Lebanon follow either the American or the European model of medical education. American-based programs consist of four years of medical studies following a bachelor’s degree, while European-based programs offer seven years of undergraduate medical education including an internship year.(12,13) The curriculum design and content also differ between the two systems.(12)

The purpose of this study was to assess knowledge of sleep medicine among final year medical students in both American and European medical education systems in Lebanon using the standardized Assessment of Sleep Knowledge in Medical Education (ASKME) survey,(14) and to determine the level of sleep knowledge among graduating medical students as they are going to be the first line of deference against worsening healthcare crisis in the country.

## Methods

### Study population

The study was conducted between January 2020 and March 2021, and the study protocol was approved by the Institutional Review Board at the Lebanese American University. The target population was final year medical students at seven Lebanese medical schools over two academic years, 2019-2020 and 2020-2021. Only final year medical students were targeted to ensure that sufficient time is provided for sleep medicine teaching throughout the medical school years, and to account for curriculum differences between schools.

The survey was disseminated electronically through email and social media platforms to all final year medical students in Lebanon, which constituted a total of 542 students for the academic year 2019-2020 and around 472 students for the academic year 2020-2021. The number of final year medical students was obtained after establishing contact with participating universities. A detailed informed consent was present at the beginning of the survey, and participants were informed that completing the survey would take around 10 minutes and the responses will remain confidential and anonymous. The faculty responsible for medical school curricula design at each of the seven universities were also contacted to fill a short survey inquiring about details regarding sleep medicine education.

### Questionnaires

A survey including questions on medical school, age, and gender of participants was administered in English due to the trilingual nature of university students in Lebanon.(15) Sleep medicine knowledge assessment was then performed using the Assessment of Sleep Knowledge in Medical Education (ASKME) survey. ASKME is a validated tool for evaluation of sleep medicine knowledge consisting of 30 questions tackling five areas of sleep knowledge: 1) basic sleep principles, 2) circadian sleep/wake control, 3) normal sleep architecture, 4) common sleep disorders, and 5) the effects of drugs and alcohol on sleep.(14) The questions are answered as “True, “False”, or “Don’t Know” to minimize random selection. On the other hand, faculty who were in charge of developing and implementing medical school curricula were surveyed on the number of hours dedicated to sleep medicine, the availability of a sleep center at the primary university hospital, and other details pertaining to sleep medicine education such as teaching methods and barriers for effective sleep medicine learning. The number of hours dedicated to sleep medicine were classified into one of three categories: 0 hours, less than 3 hours, and 3 or more hours. A cutoff of 3 hours was used for analysis of sleep medicine instruction because it is the average amount of time currently dedicated to sleep medicine education.(16,17)

### Statistical analysis

Sleep knowledge scores on the ASKME survey were transformed into and presented as proportions of correct answers by dividing the total number of correct answers by 30 (the maximum score). Data were summarized as numbers and percentages for categorical variables and mean and standard deviation for continuous ones. The association between sample characteristics and sleep knowledge scores was assessed by the Student’s t-test and Wilcoxon signed rank test if the data was not parametric. Kolmogorov-Smirnov test was used to test for a normal distribution of data. A p-value cut-off point of 0.05 at 95% confidence level was deemed statistically significant. Statistical analysis was performed using STATA 13.0.

## Results

For the class of 2020, 158 out of the 542 surveyed final year medical students responded to the survey, resulting in a 29% response rate. 80 were males (50.6%) and 78 were females (49.4%), and the majority were between 23-24 years of age (64.2%). Slightly more than half of the students were enrolled in a European-based medical education program (51.9%) while the remaining were attending an American-based program.

For the class of 2021, University 1 was not due to the lack of responses. 58 out of the 472 surveyed final year medical students responded to the survey, resulting in a 12.3% response rate. Of those, 23 were males (39.7%) and 35 were females (60.3%), and the majority were between 23-24 years of age (81.0%). More than half of the students were enrolled in a European-based medical education program (67.2%) while the remaining were attending an American-based program.

As shown in Table 1- Description of study respondents’ characteristics for 2020 and 2021, the majority of Lebanese medical schools had sleep medicine education as part of their curriculum (71.4%) and dedicated 3 or more hours for teaching sleep medicine within the curriculum (57.1%). Four out of seven medical schools had a sleep center in the primary affiliated university hospital. The Pulmonary/Critical Care specialty was the most commonly involved in teaching sleep medicine, followed by Family Medicine. The most common methods used for teaching sleep medicine were lectures and interactive discussions.

**Table 1 -.**
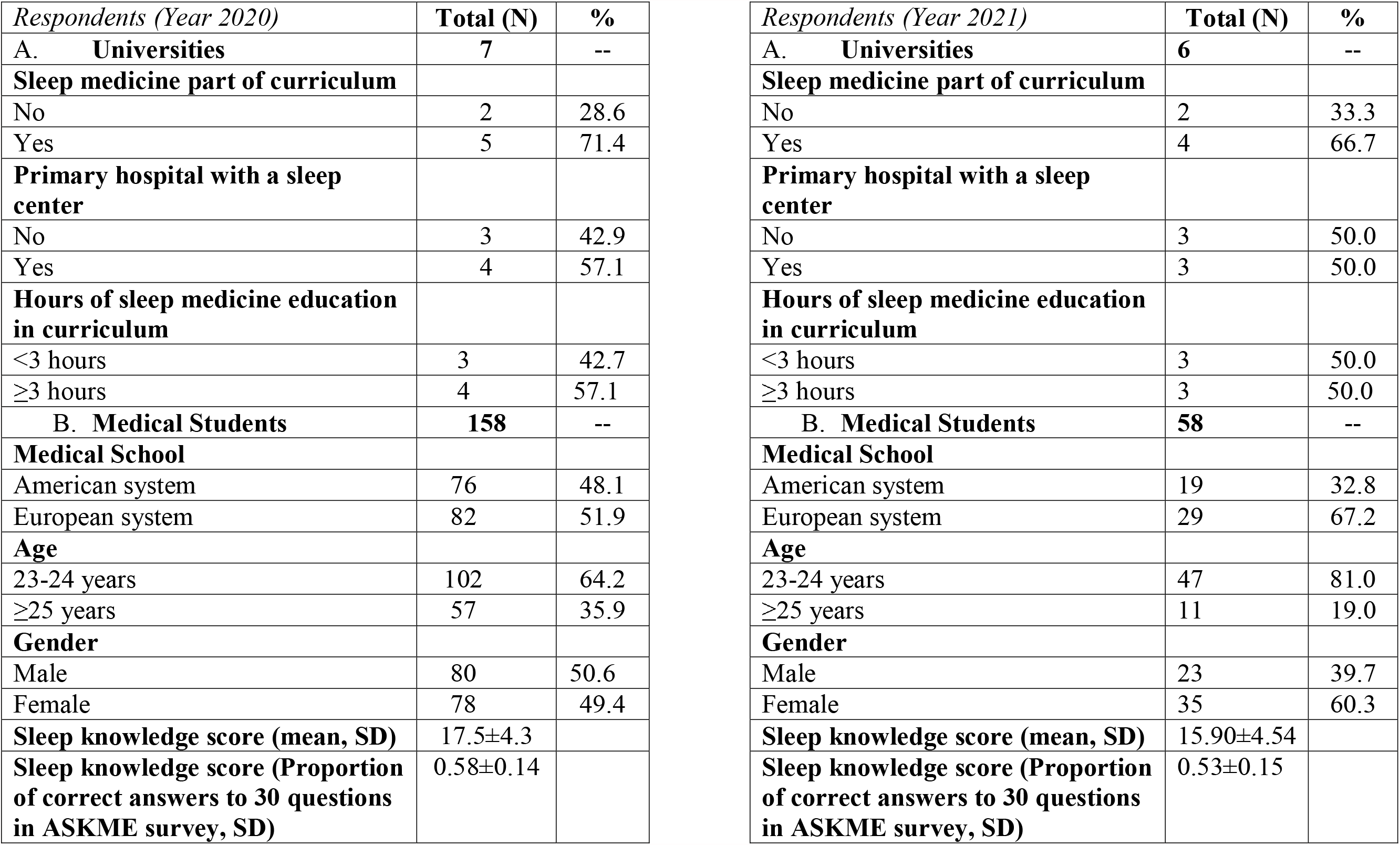
Description of study respondents’ characteristics for 2020 and 2021

| *Respondents (Year 2020)* | **Total (N)** | **%** |
| --- | --- | --- |
| **A. Universities** | **7** | **--** |
| **Sleep medicine part of curriculum** |  |  |
| No | 2 | 28.6 |
| Yes | 5 | 71.4 |
| **Primary hospital with a sleep center** |  |  |
| No | 3 | 42.9 |
| Yes | 4 | 57.1 |
| **Hours of sleep medicine education in curriculum** |  |  |
| <3 hours | 3 | 42.7 |
| ≥3 hours | 4 | 57.1 |
| **B. Medical Students** | **158** | **--** |
| **Medical School** |  |  |
| American system | 76 | 48.1 |
| European system | 82 | 51.9 |
| **Age** |  |  |
| 23-24 years | 102 | 64.2 |
| ≥25 years | 57 | 35.9 |
| **Gender** |  |  |
| Male | 80 | 50.6 |
| Female | 78 | 49.4 |
| **Sleep knowledge score (mean, SD)** | 17.5±4.3 |  |
| **Sleep knowledge score (Proportion of correct answers to 30 questions in ASKME survey, SD)** | 0.58±0.14 |  |

| *Respondents (Year 2021)* | **Total (N)** | **%** |
| --- | --- | --- |
| **A. Universities** | **6** | **--** |
| **Sleep medicine part of curriculum** |  |  |
| No | 2 | 33.3 |
| Yes | 4 | 66.7 |
| **Primary hospital with a sleep center** |  |  |
| No | 3 | 50.0 |
| Yes | 3 | 50.0 |
| **Hours of sleep medicine education in curriculum** |  |  |
| <3 hours | 3 | 50.0 |
| ≥3 hours | 3 | 50.0 |
| **B. Medical Students** | **58** | **--** |
| **Medical School** |  |  |
| American system | 19 | 32.8 |
| European system | 29 | 67.2 |
| **Age** |  |  |
| 23-24 years | 47 | 81.0 |
| ≥25 years | 11 | 19.0 |
| **Gender** |  |  |
| Male | 23 | 39.7 |
| Female | 35 | 60.3 |
| **Sleep knowledge score (mean, SD)** | 15.90±4.54 |  |
| **Sleep knowledge score (Proportion of correct answers to 30 questions in ASKME survey, SD)** | 0.53±0.15 |  |

The mean overall score on sleep medicine knowledge was 17.5/30 for the year 2020 and 15.9/30 for the year 2021. Further analysis showed that 24.7% of participating medical students scored above 21/30, 71.5% scored between 10 and 20, and 3.8% scored less than 10/30 for the year 2020. Whereas for class of 2021, analysis showed that 13.8% of participating medical students scored above 21/30, 79.1% scored between 10 and 20, and 5.1 % scored less than 10/30.

As shown in Table 2- Proportion correct on ASKME survey according to sleep medicine education hours per university, years 2020, 2021, the score proportions for the medical schools for the year 2020 ranged between 0.57 and 0.59 with two exceptions: University 6, which had no sleep medicine education in the curriculum, had a score proportion of 0.71, and University 7, which has more than 3 hours of sleep medicine education had a score proportion of 0.52. However, for the 2021 cohort, the score proportions ranged between 0.52 and 0.58 with one exception being University 4.

**Table 2 -.** Proportion correct on ASKME survey according to sleep medicine education hours per university, years 2020, 2021

|  | **Sleep medicine education hours in curriculum**<br>**(2020)** |  |  |
| --- | --- | --- | --- |
| **University** | **0 hrs** | **<3 hrs** | **≥3 hrs** |
| 1 | -- | -- | 0.59 |
| 2 | -- | -- | 0.58 |
| 3 | -- | 0.58 | -- |
| 4 | -- | -- | 0.57 |
| 5 | 0.57 | -- | -- |
| 6 | 0.71 | -- | -- |
| 7 | -- | -- | 0.52 |

|  | **Sleep medicine education hours in curriculum**<br>**(2021)** |  |  |
| --- | --- | --- | --- |
| **University** | **0 hrs** | **<3 hrs** | **≥3 hrs** |
| 1 | -- | -- | 0.0 |
| 2 | -- | -- | 0.53 |
| 3 | -- | 0.53 | -- |
| 4 | -- | -- | 0.41 |
| 5 | 0.58 | -- | -- |
| 6 | 0.52 | -- | -- |
| 7 | -- | -- | 0.56 |

Based on students’ responses, the true-or-false questions with the highest percent of correct answers were: “Melatonin is a natural body hormone that typically increases during nighttime hours,” and “Weight loss is often indicated in the treatment of primary snoring or mild obstructive sleep apnea.”, while the question with the lowest percent of correct answers was, by far: “In alcoholics in recovery, sleep normalizes within one month of alcohol abstention” for both years.

All groups had similar sleep knowledge scores; there was no difference in mean knowledge scores between gender, age, and medical school system. Also, there was no significant difference in mean scores on sleep medicine knowledge between medical schools that included sleep medicine in their curriculum versus medical schools that did not include sleep medicine education. No significant difference in sleep knowledge scores was found between medical schools with less than 3 hours and 3 or more hours of dedicated sleep medicine education in the curriculum for both years. Also, no significant difference in sleep knowledge scores was seen between medical schools affiliated with a primary hospital or not. Over 40% of the medical schools saw no obstacle in implementing sleep medicine education within their curriculum as seen in

Figure 1, while over a quarter replied that lack of sufficient time in the curriculum was a major obstacle in implementing sleep medicine education in their curriculum. The remainder of responses were divided by a perceived lack of interest in the subject matter and lack of resources and expertise in sleep medicine.

**Figure 1 -.**
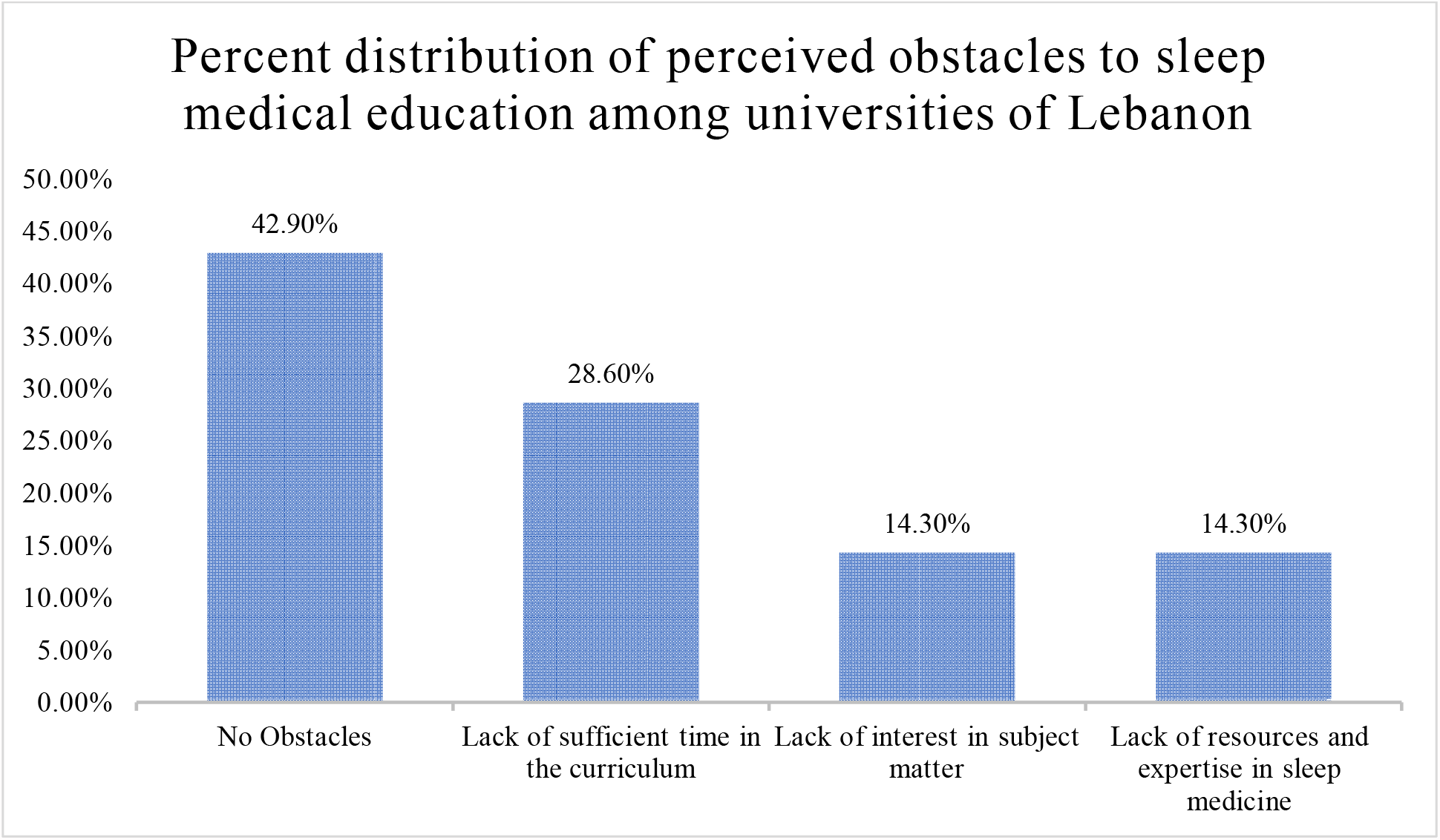
Percentage of perceived obstacles to sleep medicine education

## Discussion

The aims of the study were to assess the level of sleep knowledge among final year medical students in Lebanon as a surrogate marker of their level or preparedness as they enter healthcare in a country in economic and political turmoil. It also aims to look at the association between said knowledge and presence or absence of formal sleep education in their respective curricula. The mean overall score on ASKME in our cohort was 17.5±4.3 out of 30 for 2020 and 15.9±4.5 out of 30 for 2021. This study demonstrated that higher scores on ASKME were linked with having sleep medicine education in the medical school curriculum. It suggests that incorporating sleep medicine education in the curriculum may improve understanding of sleep principles and pathologies. In addition, it highlights the readiness of graduating students to tackle sleep health issues in a country of crisis.

The average knowledge in sleep medicine among final year medical students in Lebanon was relatively better than other regions outside the US. The mean proportion of knowledge scores in this study was found to be 0.58 ± 0.14 and 0.53±0.15 for 2020 and 2021, respectively, which is higher than the mean proportion of scores for students in Croatia (mean proportion = 0.41)(18) and Saudi Arabia (mean proportion = 0.35)(17). Also, in Singapore, the mean proportion was found to be 0.41 for 5^th^ year medical students.(19) In fact, 24.7% and 13.8% of Lebanese medical students scored above 21/30 on the ASKME survey, whereas only 2.5% of participating medical students in Egypt, 1.7% in Singapore, and 0.6% in Saudi Arabia scored between 21 and 30 points on the same questionnaire.(17,19,20) The high average knowledge among final year medical students in Lebanon is particularly important during the ongoing Lebanese crisis. The high sleep literacy among graduating physicians in both years ensures that the graduating classes are well equipped to face a surge of sleep disturbances originating from decreased quality of life and social uncertainty caused by the economic and political collapse. Studies have linked poverty with poor sleep and poor health and quality of life.(4) The proportion of people living in poverty has exponentially increased in Lebanon since late 2019, resulting in brief bursts of violence culminating in the cataclysmic explosion in Beirut port.(1,21) This has particularly had a negative impact on sleep and health.(22)

It is important to note that most Lebanese medical schools use the National Board of Medical Examiners (NBME) tests to evaluate students at the end of each module and clerkship, and these exams tackle sleep disorders within the “Nervous System and Special Senses” section.(23) Moreover, there is a high trend among Lebanese medical students to pursue residency training in Europe or in the US. In fact, since 1978, the number of Lebanese medical school graduates that train in the US has been trending upwards at a rate of about 71 additional graduates per year.(24) Therefore, some students spend a significant amount of time preparing for standardized exams like the United States Medical Licensing Examination (USMLE) which could make them more knowledgeable in fields that are not adequately covered within their medical school curricula. It is important to note, however, that given the overall low number of hours currently dedicated to sleep medicine education in Lebanese medical schools’ curricula, sleep medicine knowledge scores may improve with greater incorporation of sleep medicine topics in the syllabus.

The questions with the highest and lowest number of correct answers on the ASKME survey in this study were similar to those reported by Almohaya et al.(17) In general, questions tackling basic sleep principles and circadian sleep/wake control had higher percentages of correct answers than questions tackling sleep architecture as well as the effects of drugs and alcohol on sleep. For example, only 13.3% of respondents in 2020, 6.9% in 2021 in this study and 14.9% in the Saudi study knew that sleep does not normalize in recovering alcoholics within one month of abstention.(17) This suggests that medical students may be familiar with the basics of sleep medicine but not the impact of recreational substances on sleep.

Four out of seven medical schools in Lebanon spend 3 hours or more on sleep medicine education. The hours dedicated to sleep medicine education in Lebanon are comparable to countries like the US and Australia.(25) In Saudi Arabia, on the other hand, the average amount of time spent on sleep education was just under 2.5 hours, and Malaysia, Indonesia, and Vietnam provide no sleep education.(25) Despite Lebanese medical curricula’s higher incorporation of sleep medicine, the Middle East still lacks adequate exposure to sleep medicine during medical training, and the majority of universities lack postgraduate sleep medicine fellowships.(20) It is essential to provide sleep medicine education through an integrated experience of didactics and clinical, hands-on exposure for a more comprehensive understanding of the field among medical students. In fact, The Institute of Medicine (IOM) report recommends that sleep medicine exposure should begin prior to entering into residency and early on as part of the medical school curricula.(26)

Despite the high prevalence of sleep disorders and their effect on health, 53% of curriculum organizers across several countries still believe that teaching sleep medicine should be a low priority for medical students.(25) The main obstacle to implementing adequate sleep medicine education in the curriculum in Lebanese medical schools is curriculum overload. Time constraints have similarly been considered a major obstacle in Saudi Arabia and across 12 other countries according to Mindell et al.(17,25) It is understandable that medical schools must address a substantial amount of scientific and clinical material in their curricula in a limited time, making it difficult to devote a block for sleep and sleep disorder education. However, allowing students to take elective courses in sleep medicine clinics, integrating sleep history and physical signs and examination into clinical medicine, and using computer-based simulations to expose students to sleep disorders are some of the solutions that have been proposed to tackle the gap in sleep medicine proficiencies.(27,28)

Of the limitations of our study is, first and foremost, the low response rate in both years which, nevertheless, was similar to that of other studies.(29) The low response rate of our study precludes generalizability to our target population. The constant stress of living with political instability and economic collapse may have affected the motivation of the potential responders. However, it has been shown that the response rate for email surveys is usually lower than traditional surveys, between 25% and 30%.(30) We also do not have data about the non-responders, who may differ from responders in terms of characteristics such as GPA, future specializations, interests, and standardized exam preparations. Another limitation is the lack of information on time elapsed since responders was exposed sleep medicine curriculum at his or her institution, which may influence knowledge retention and thus our study results. This was, however, a descriptive pilot study to gather data that may be used for future, more rigorous, educational research in this field. Strengths of the study include the use of a validated tool (ASKME) for the assessment of sleep medicine knowledge, and the inclusion of participants from both the American and European systems of medical education; moreover, the faculty responsible for organizing the curriculum at all seven universities were contacted and surveyed to receive accurate and detailed information about sleep medicine education at their respective institutions. To our knowledge, this is the first study in Lebanon to explore sleep medicine education among final year medical students.

## Conclusion

To conclude, incorporating sleep medicine education in the medical school curriculum was shown to correlate with higher scores on the standardized ASKME survey. Medical students in Lebanon possess good knowledge in sleep medicine relative to other Arab and European countries. Nonetheless, more effort should be made in incorporating sleep medicine in all medical school curricula and in increasing clinical exposure. This study shed an international perspective on the subject of sleep medicine education. Future research should further explore the effect of increasing sleep medicine clinical exposure on knowledge and prompt recognition of sleep disorders.

## Abbreviations

ASKME: Assessment of Sleep Knowledge in Medical Education
IOM: The Institute of Medicine
NBME: National Board of Medical Examiners
US: United States
USMLE: United States Medical Licensing Examination

## Declarations

### Data Availability

Data available on request due to privacy/ ethical restrictions

### Ethics approval

The study was performed in accordance with the Declaration of Helsinki and was approved for an exemption by the Institutional Review Board at the Lebanese American University (RE#: LAU.SOM.SB1.16/Jan/2020)

### Consent for publication

Not Applicable

### Author contributions

**R.A**.: Methodology, Investigation, Writing – Original Draft. **C.C**.: Formal Analysis, Writing-Original Draft. Writing-Review & Editing. **V.Z**.: Formal Analysis, Data Curation, Writing, Reviewing, Editing **H. A**.: Conceptualization, Methodology, Writing-Review & Editing. **S.A.B**.: Conceptualization, Investigation, Writing-Review & Editing. All authors read and approved the final manuscript.

## Acknowledgement

Not Applicable

